# EsViritu is a sensitive and specific profiler for viral pathogen detection and genome recovery in metagenomes

**DOI:** 10.64898/2026.09.29.755457

**Authors:** Michael J. Tisza, Shelby R. Simar, Jackie K. Zorz, Sara J. Javornik Cregeen, Anthony W. Maresso

## Abstract

Metagenomic sequencing allows for detection of any number of pathogenic viruses from a single sample without a priori knowledge of its contents. This has created a substantial opportunity to characterize pathogens in humans and their environment, but the field is held back by a lack of bioinformatics tools for taxonomic profiling and reconstructing virus genomes. The ideal profiler would simultaneously do two things: report viruses at single-read sensitivity so that ultra-low-abundance pathogens are not missed and hold near-perfect specificity so that a pathogen is never falsely reported. Here we present EsViritu, a taxonomic profiler that pairs an iterative greedy clustering algorithm and lowest-common-ancestor taxonomic refinement with a curated Virus Pathogen Database to reconstruct virus genomes and generate virome profiles directly from reads. We benchmarked EsViritu against four leading metagenomic profilers (Kraken2, Centrifuger, Metabuli, and Sylph), running each program across a full parameter sweep and against the same Virus Pathogen Database for an apples-to-apples comparison. Across synthetic positive controls, pathogen-free negative controls, and mutated mock communities, EsViritu was the most accurate tool at reconstructing community composition: it led or tied every tool on relative abundance accuracy at all taxonomic ranks and led or tied on detection across ranks. Importantly, other tools frequently report the presence of multiple related taxa when only reads for a single genome are present, but EsViritu faithfully records the correct, single taxon and genome. EsViritu achieves read-level sensitivity for sequences as low as 85% average nucleotide identity to the nearest reference and reports essentially no false positives in pathogen-free host and synthetic samples. EsViritu also returns reconstructed consensus genomes whose nucleotide identity to the nearest reference closely tracks ground truth, yielding both a profile and genomes from a single run.

## Introduction

Virus metagenomics is rapidly becoming an essential strategy for unbiased pathogen detection^1–4^. Where clinical and surveillance pipelines once targeted a single virus at a time, untargeted sequencing can contain sequences from whole viral communities from a single specimen, including unexpected pathogens. The applications span public health surveillance, biosecurity, and the investigation of diseases of unknown etiology^5–7^. The possible sample types are similarly diverse, including clinical specimens, wastewater, air filters, and a wide range of other environmental matrices^8–11^. Hybrid-capture enrichment and ultra-deep sequencing of virus-like-particle preparations now routinely deliver the read depth needed to find rare viruses^8,12,13^, which shifts the bottleneck from the bench to the analysis.

The bioinformatics workflow is a rate-limiting step that determines the fidelity of surveillance efforts, yet the current landscape of profilers presents important tradeoffs^14,15^. Some pipelines are ultra-heavy, aligning every read against all of NCBI, which is costly, slow, and difficult to adopt for most labs^16–18^. Other tools were built for a single study and are no longer maintained. Genomics scientists therefore often use general-purpose taxonomic profilers^19^, but those tools are tuned for bacterial communities or for whole-NCBI searches^20^, and their assumptions do not hold for viral genomes, which are short, fast-evolving, and span enormous diversity. At the same time, most tools are designed to return a taxonomic profile and nothing more. However, virus surveillance needs both a taxonomic profile and genome-level reconstructions to confirm a call and to track strain-level variation. These limitations of genomic algorithms are costly: missing an ultra-low-abundance pathogen could mean a missed outbreak, while falsely reporting a pathogen that is not present can trigger an unwarranted and expensive public-health response.

*EsViritu* is a sensitive and accurate profiler built specifically for viral pathogens. It combines single-read-level sensitivity with state-of-the-art specificity through three design choices. First, an iterative greedy clustering algorithm selects the single best reference genome from a set of related strains, so reads likely derived from a single strain are consolidated onto one representative rather than scattered across many near-identical references^21^ (Fig. 1A). Second, taxonomic assignments are refined by lowest-common-ancestor placement, which provides confident taxonomic calls to ranks supported by the data^22^. Third, the entire workflow is (optionally) anchored to the Virus Pathogen Database (Fig. 1B)^23^, curated by the authors, which increases both speed and specificity by collapsing near-identical genomes^24^, removing contaminated or problematic references, and avoiding alignment to large non-viral genomes^25^. Still, this database can be replaced or customized easily with built-in *EsViritu* commands. *EsViritu* produces both a taxonomic profile and a set of consensus genome reconstructions, each annotated with average nucleotide identity (ANI) to the nearest reference (Fig. 1A). A previous version of *EsViritu* was first used in Tisza et al. 2023^8^, but the algorithm, codebase, and database have been rebuilt from scratch to increase specificity and sensitivity.

**Figure 1.**
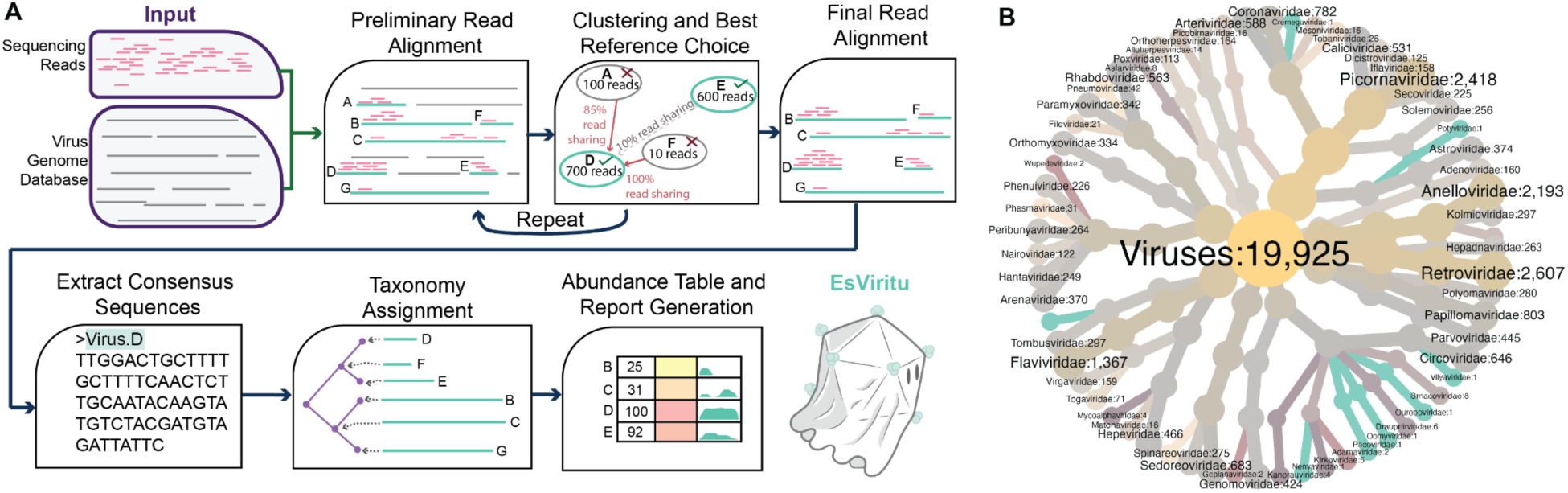
EsViritu and Virus Pathogen Database Overview. (A) The inputs for *EsViritu* are sequencing reads and the Virus Pathogen Database. After a preliminary alignment, an iterative greedy clustering step resolves read sharing among related references and selects the best genome from each cluster, repeating once to decide the final reference set. Reads are then realigned to the chosen references, consensus sequences are extracted, taxonomy is assigned by lowest-common-ancestor placement, and an abundance table and report are generated. (B) The Virus Pathogen Database is composed of sequences and their metadata derived from 63 viral families of putative pathogenicity to humans, animals, and plants (see Methods).

Here we benchmark *EsViritu* against the leading metagenomic profilers and show that it is the most accurate tool for reconstructing viral metagenomes. We evaluated five tools across real virus genomes as positive controls spanning 76% to 95% ANI to their closest database reference, pathogen-free negative controls drawn from a variety of environmental and animal backgrounds and mutated mock communities with realistic abundance structure. To ensure an equal comparison, each tool was run across a full parameter sweep and queried against the same *EsViritu* Virus Pathogen Database version. *EsViritu* avoids false positives across diverse background samples, reaches read-level sensitivity for sequences as low as 85% ANI to reference, resolves taxonomic profiles, and is the most accurate tool for recovering relative abundance. EsViritu is available on <u>Bioconda</u> and the source code is on GitHub at <u>cmmr/EsViritu</u>, with full documentation on a <u>readthedocs</u> website.

## Results

All five tools were evaluated against the same database, and each tool was tested with a range of parameters that govern its sensitivity and specificity. For *EsViritu* short-read “*sr*” (conservative) and “*sense*” sensitive read alignment seeding parameters were tested. For *Kraken2*^19^, we tested confidence score (0 – 0.5) and minimum-hit-group (number of overlapping k-mers required per call) (2 – 5) thresholds. For *Centrifuger*^26^, minimum hit length (auto, 50, 100) and alignment score (0 – 500) were tested. For *Metabuli*^27^, we varied precision threshold (0, 1) and minimum-score settings (0, 0.15). And, for *Sylph*^28^, minimum k-mer count per genome was tested (10, 20, 50). All headline comparisons use each tool’s best-performing parameter set, which allays any concern that a tool was impaired by suboptimal settings. Every tool was queried against the same version of the *EsViritu* Virus Pathogen Database, so differences in performance reflected the intrinsic algorithms rather than differences in reference content.

### *EsViritu* reaches read-level sensitivity while avoiding false positives

We first compared the minimum number of reads each tool needs to detect a virus, and how that sensitivity degrades as the virus genome diverges from its nearest reference. Synthetic read sets were generated from seven real virus reference genomes spanning 75.9% to 95.6% ANI to their closest database reference, at sequencing depths of 1, 10, 100, and 1,000 reads, with 10 independent permutations at each depth. Detection probability rose with depth for all tools, and the tool sensitivity varied substantially at the low-depth end where it matters most for rare pathogen detection (Fig. 2A). Further, decreasing ANI affected the tools’ ability to detect viral reads, and were parameter-dependent for all tools, especially *Kraken2,* and *Centrifuger* (Fig. 2A). *EsViritu* using *sr* and *sense* settings was near the top, with single-read sensitivity for genomes in 85%-96% ANI range and still detecting genomes of 75%-76% ANI with 10 or more reads. *Metabuli*, with its translated search approach, was the most sensitive regardless of parameters.

**Figure 2.**
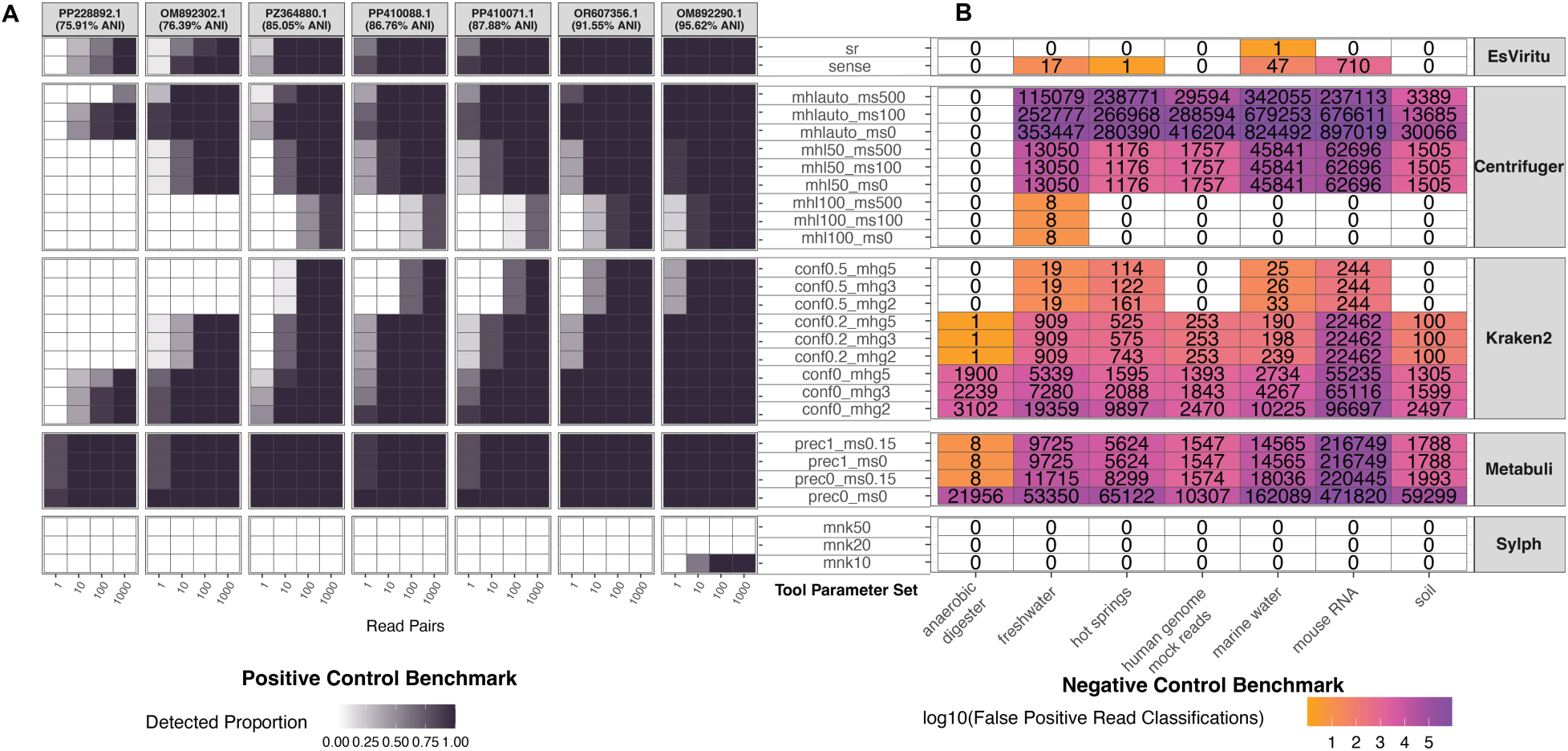
Precision in Positive and Negative Controls. (A) Detection probability versus sequencing depth for each tool and parameter set for reads from seven real virus genomes of varying divergence from closest reference. (B) False positive read assignments for each tool and parameter set across seven qualitatively different sequencing sets including both environmental metagenomes and vertebrate genomes.

A profiler is only useful if it reports nothing when no pathogen is present, so we ran all five tools on seven pathogen-free samples spanning host, synthetic, and environmental backgrounds (human genome mock reads, lab mouse RNA, anaerobic digester, freshwater, marine water, hot springs, and soil) and counted false-positive reads and false species (Fig. 2B, Fig. S1C, Table S6, S16-S20). *EsViritu* (*sr* mode) and *Sylph* (all parameters) were effectively clean, each with a median of zero false species and *EsViritu* with one spurious call across all samples. *EsViritu* with more permissive settings (*sense* mode) showed between 0 and several hundred false-positive reads.

For the other tools, the parameters that allowed them to perform well at the positive control benchmark also caused them to call a very high number of false positive reads and species (Fig. 2B, Fig. S1C). For instance, *Centrifuger* showed dramatic differences depending on parameter choice, going from near-0 false positive reads when using *minimum hit length=100*, but becoming unspecific in more permissive settings (hundreds of thousands of false hits/read library). *Kraken2* had similar parameter-dependence, carrying a small but nonzero false positive rate (median 6 false species, maximum 24; 0 to 244 false positive reads) with confidence=0.5, but failing to control false positives at confidence=0.2 or 0^29^. *Metabuli* was the clear outlier, reporting a median of 18 false species and as many as 71 in a single sample for its *most* conservative setting (8 to 216,749 false positive reads) (Fig. S1C). The differences across tools were significant for both false-positive read percentage and false-species count (Friedman p = 9.8 x 10^−5^ and p = 4.2 x 10^−4^, Table S7), and *Metabuli*’s false-positive read percentage was significantly higher than that of *EsViritu*, *Centrifuger*, *Kraken2*, and *Sylph* in post-hoc tests (Benjamini-Hochberg adjusted p = 0.039) (Table S8). These benchmarks allowed us to generate “headline” parameters for each tool that maximized specificity but retained as much sensitivity as possible (see Methods) (*EsViritu=sr; Centrifuger=minimum-hit-length:100, minimum score:0; Kraken2=confidence score:0.5, minimum-hit-group:2; Metabuli=precision:1, minimum score:0.15; Sylph=10 k-mers-per-genome*).

We next summarized each genome by its LD50, the read depth at which a tool reaches 50% detection probability and compared it to genome ANI (Fig. S1A, Tables S4, S5). *Metabuli* reached 50% detection at the lowest depths across the panel, with most LD50 values at or near a single read. *EsViritu* held read-level sensitivity (LD50 near one read) down to 85.05% ANI and required only a handful of reads at the two most divergent genomes (75.9% and 76.4% ANI). *Kraken2* and *Centrifuger* with these settings did not detect genomes below 85% ANI and did not reach single read sensitivity until 95% ANI. *Sylph*, by contrast, did not reach 50% detection in six of the seven genomes at any tested depth.

To determine probability of detection in varying conditions, we used a Firth-penalized logistic regression of detection on tool, depth, and genome. The model-estimated detection probability for *EsViritu* was 0.995. Only *Metabuli* detected more often (odds ratio 127.6 versus *EsViritu*, p = 0.012; estimated detection probability 0.99996), a result best read alongside its tendency to over-call seen in the negative controls below. For headline parameters, every other tool detected dramatically less often than *EsViritu*: *Kraken2* (odds ratio 1.3 x 10^−3^, p = 1.3 x 10^−23^), *Centrifuger* (odds ratio 2.7 x 10^−4^, p = 4.0 x 10^−28^), and *Sylph* (odds ratio 6.4 x 10^−8^, p = 2.4 x 10^−9^; estimated detection probability 1.8 x 10^−3^) (Table S1-S3). Because ANI is confounded with genome identity in this design (one genome per ANI level), divergence is shown descriptively rather than as a fitted coefficient. Some *Kraken2* and *Centrifuger* parameter combinations allow detection of more divergent genomes (Fig. S1B), but these parameters have high false positive rates (Fig. 2B).

**Figure S1.**
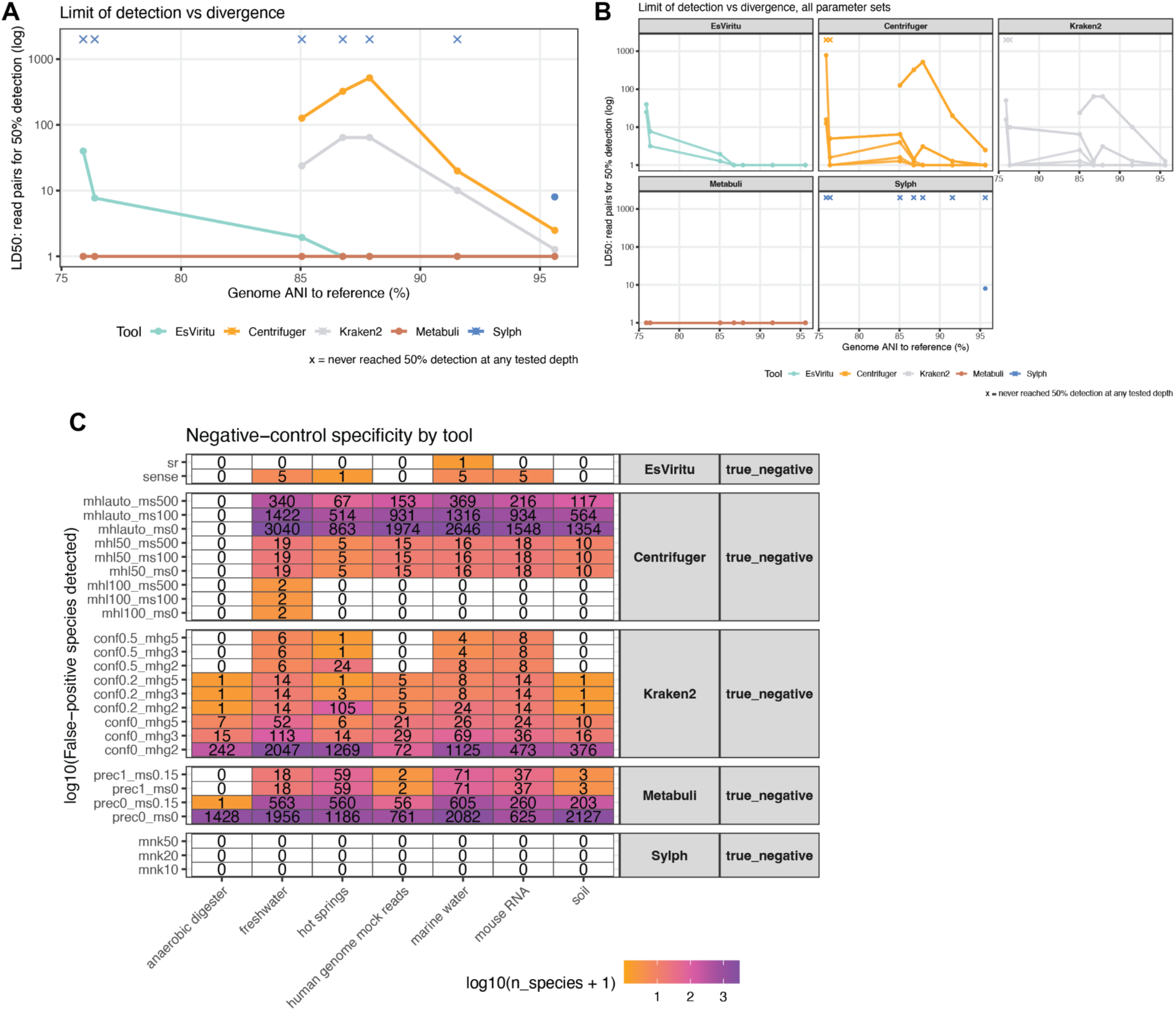
Additional Positive and Negative Control Analyses. (A) Limit of detection for positive controls by ANI (x-axis) and estimated LD50 (y-axis) for “best” parameter by tool. (B) Like (A) but showing all parameters and tools. (C) Negative control libraries across tools and parameters, measured for number of unique species reported.

### *EsViritu* is an accurate profiler of mock virus communities

To test whole-community accuracy with a known ground truth, we built mutated mock communities at two levels of complexity. Three “random” mock communities each contained 90 viral genomes drawn randomly from the Virus Pathogen Database, and three further “one-per-family” mock communities each contained 60 genomes drawn so that no more than one genome came from any family, to measure intra-family specificity (Fig. 3A). In every community the genomes were mutated with *simuG*^30^ to a controlled ladder from 100% (no mutation) down to 80% ANI to their references; genome-level abundances were drawn from a log-normal distribution. Then, an extensive “background” set of reads was added synthesized from 100 random bacterial genomes (from NCBI) plus the human genome. The background set models the complexity of real metagenomic and environmental samples. Synthetic read pairs were generated with InSilicoSeq to 10,000 total viral reads and 50,000,000 background reads^31^.

**Figure 3.**
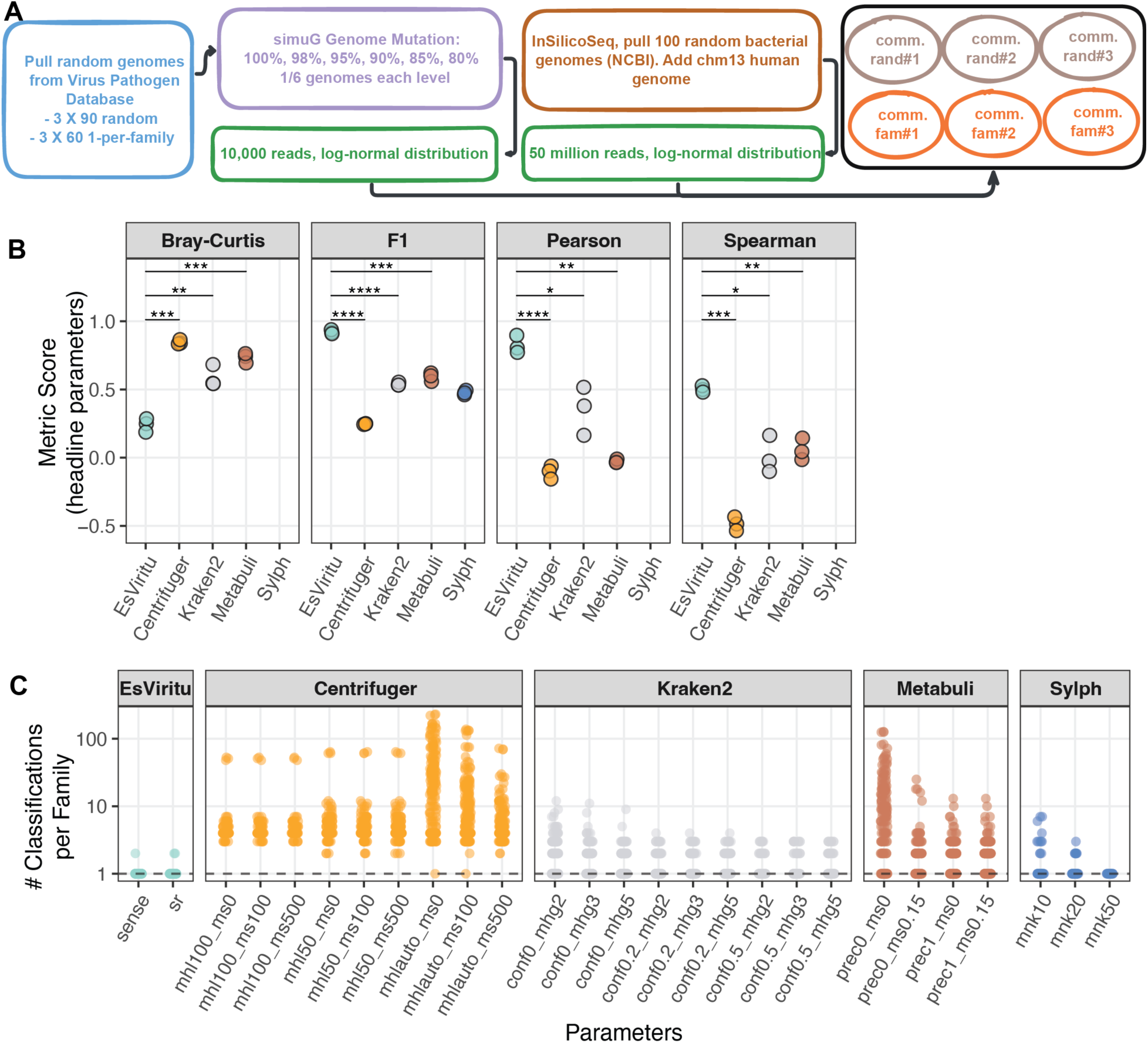
*EsViritu* Excels at Reconstruction Accuracy of Mock Communities. (A) Mock virome community generation workflow for six distinct communities with varying mutation, abundance, and taxonomic composition. (B) Using one-per-family mock communities and “*divergence-capped*” metrics, each tool’s “headline” (best) setting was compared for Bray-Curtis Dissimilarity (lower=better), F1 score (higher=better), Pearson and Spearman correlation (higher=better) *EsViritu* performs better than all other tools in these benchmarks. Each panel uses student’s t-test for comparisons. (C) Using one-per-family mock communities, when only one taxon should be classified per-family, each tool and parameter is measured as number of classifications per-family. Only *EsViritu* properly consolidates taxonomic profiles properly.

**Figure S2.**
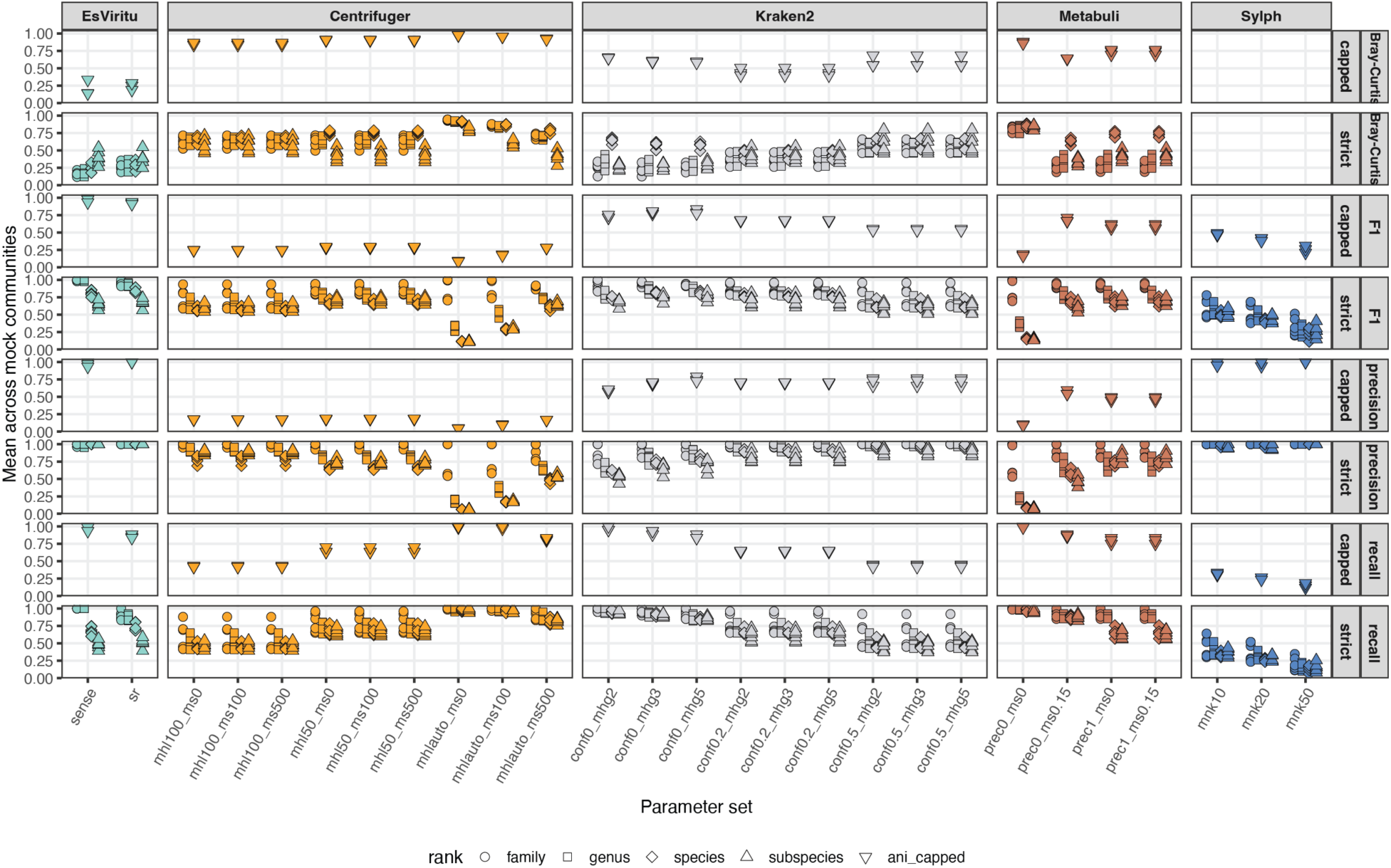
Mock Community Analysis Parameter Sweep. Each tool across all parameter sets and mock communities (i.e. “random” and “one-per-family”) were run. *Divergence-Capped* metrics are calculated on an ANI-justified basis. *Strict* metrics are calculated for match at the family, genus, species, and subspecies level, regardless of ANI similarity.

Virus read classification accuracy is affected by the interaction of rank and genome divergence. E.g. we can say intuitively that reads from a genome with 80% ANI to genomes from closest known species should not be classified as that species but instead be unclassified at the species level. However, there is no standard method to score genomes with varying levels of divergence from known genomes, and viral species thresholds are family-dependent^32^. Therefore, we compared tool-based virome profiles to ground truth under two truth definitions. First, *divergence-capped* modifies ground truth taxonomy calls from divergent genomes to be unclassified below the rank its ANI-to-reference can support (subspecies: 100%, 98%, species: 95%, 90%, genus: 85%, 80%; see Methods). This capping may be reasonable, but it is also arbitrary. Second, *strict* keeps ground truth taxonomy calls the same as the reference lineage, regardless of ANI divergence. Notably, due to difficulty modeling true abundances in the “random” communities, only the “one-per-family” mock communities were measured in the *divergence-capped* metrics.

*EsViritu*, regardless of parameter choice and ground truth definition, performed better than or as well as any other tool on virtually all communities, ranks, and metrics (Fig. 3B, Fig. S2, Table S9-S12, S21-S24). We report the headline parameter for each tool with *divergence-capped* scores in Figure 3B, as these probably best represent the cleanest source of ground truth. These metrics measure both detection (F1, precision, recall) and abundance (Bray-Curtis dissimilarity, Pearson, and Spearman correlation)^33^, and *EsViritu* is significantly better for all metrics. We report both *strict* and *divergence-capped* metrics for all six communities and all tool/parameter combinations in Fig. S2, finding that *EsViritu* is robust to these permutations. *Sylph* reports a k-mer-containment sequence-abundance value and was not readily comparable to other abundance metrics.

Community profiling accuracy is affected by erroneous multi-classification of single-origin reads. Using the “one-per-family” mock community, we know that only one taxon should be reported per family. *EsViritu* achieves this, reporting a single record in its virome profile for nearly every genome in the mock community (Fig. 3C), whereas all other tools report multiple taxa for many or most sequences from a single genome. The practical implication is that *EsViritu* gives scientists and public health officials a cleaner report to interpret and better data to use for downstream studies.

Pearson correlation between tool calls and ground truth abundance (reads per million) was measured. Pearson correlation was high for all tools for 100% ANI (unmutated) genomes down to 95% ANI, with *EsViritu* being the most correlated. Most tools struggle with low (90%-80%) ANI genomes, with the exception of *EsViritu* in *sense* mode and some configurations of *Metabuli* (Fig. S3A). *EsViritu* sense mode is the only tool/parameter combination retaining a Pearson correlation above 0.75 at all ANI levels (Fig. S3A).

*EsViritu*’s performance is also reflected in the ability to detect genomes even at low ANI levels (Fig. S3B) as shown orthogonally in Figure 2A. Comparing assigned vs expected read counts for each taxa, we see that *EsViritu* in *sense* mode can quantitatively track highly divergent genomes (Fig. S3C). In addition, using Lin’s CCC, we see that *EsViritu* led in abundance metrics vs ground truth, and was nearly perfect by this measure for “resolvable” (>=95% ANI) genomes (Fig. S3D, Table S13).

**Figure S3.**
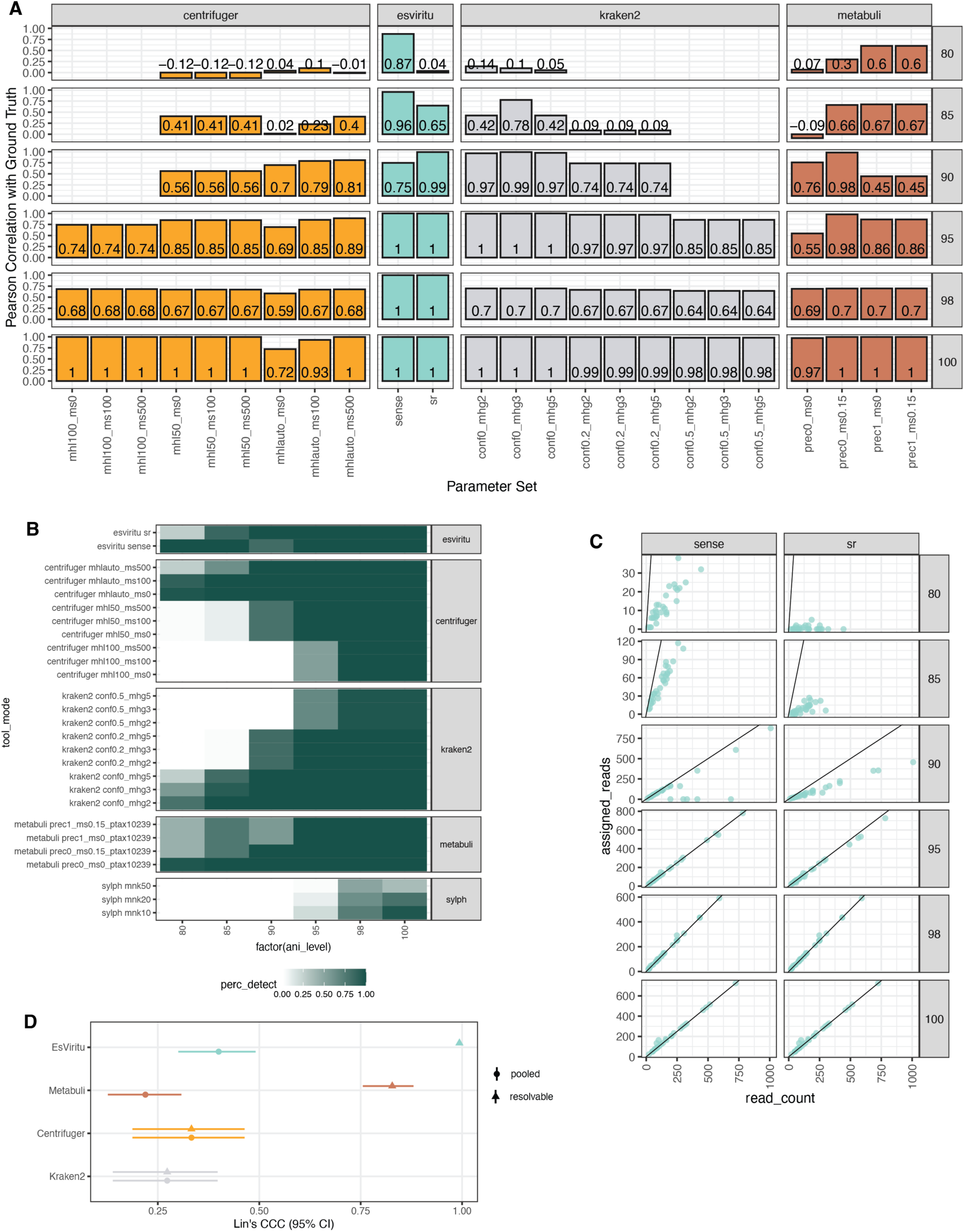
Detection and Abundance Metrics for Mock Communities. (A) Pearson correlation for measured vs. expected abundance (reads per million) for viruses at different levels of mutation (100% ANI to 80% ANI) in mock communities. (B) Detection probabilities for each tool/parameter combination for viruses at mutational levels. (C) measured vs. expected read assignment counts for EsViritu parameters (*sr* and *sense*) for genomes at different levels of mutation. (D) Lin’s CCC for all “pooled” taxa and “resolvable” (1 genome per family and >= 95% ANI).

**Figure S4.**
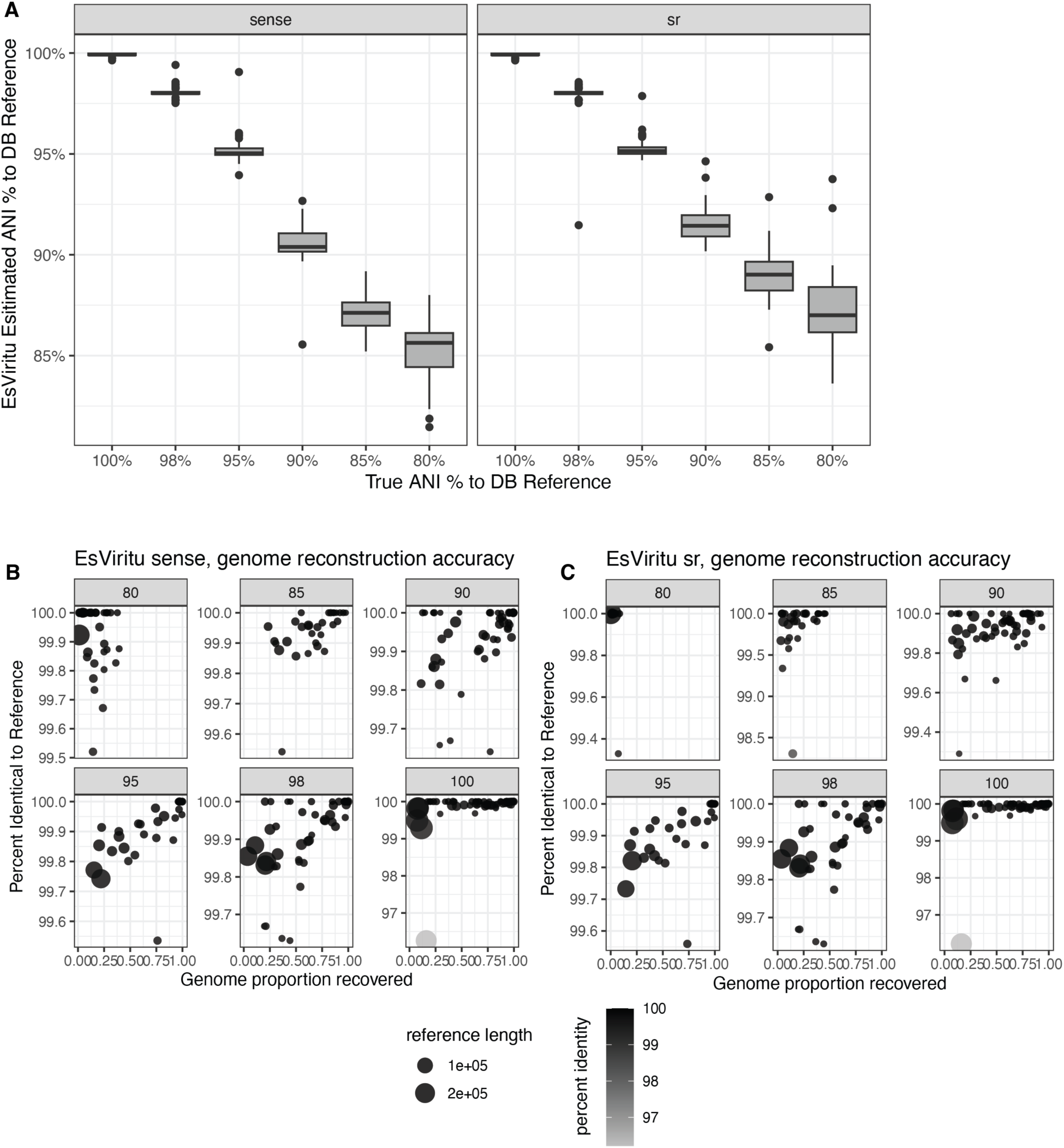
EsViritu Genome Reconstruction. (A) Estimated ANI of *EsViritu* consensus reconstructions versus ground-truth ANI across the mock-community divergence ladder; concordance is high at 90% ANI and above. (B) Comparison of EsViritu reconstructions directly with *simuG*-mutated references using BLASTN, for *sense* mode. Facets are ANI levels for mutated genomes. (C) like (B) but for *sr* mode.

### *EsViritu* Reconstructs Genomes with High Accuracy

In addition to a taxonomic table, *EsViritu* returns a consensus genome for each detection together with its ANI to the nearest reference, a feature lacking in the other profilers in this benchmark. On the mock communities, the ANI that *EsViritu* estimated for each reconstructed genome showed high concordance with the ground-truth ANI for genomes at 90% identity and above, the range where a single-genome reconstruction is well defined (Fig. S4A). Reads below 90% ANI sometimes fail to seed on the reference genome, leading to dropout of less similar reads. Further, aligning *EsViritu* reconstructions with the actual *simuG* mutated genomes (via BLASTN), we see that the results are identical or near identical for almost all reconstructions (Fig. S4B-C). The trend toward higher mismatches in genomes with low coverage suggests that the read-level technical errors can propagate to the reconstructions, probably only effecting regions with one or two reads covering. The genome reconstruction feature lets an analyst not only name a virus but place it relative to known strains, distinguishing a textbook match from a novel divergent lineage directly from the report.

### *EsViritu* scales with computation resources but has higher wall time than other tools

To assess the computational performance of each tool on real data, we processed an ultra-deep wastewater virome dataset subset from 1 million to 200 million read pairs and varied CPU availability from 2 to 32 (Table S15). Then, we measured wall time and maximum memory usage. Most tools, including EsViritu scale, if imperfectly, with CPU availability. *Sylph* does not, but it is capable of horizontal scaling by sketching multiple sequencing libraries in parallel (which was not tested). *EsViritu* typically took 2 - 8 times longer than the other programs across library size and CPU availability (Fig. 4A). Memory usage was moderate for EsViritu, whereas *Centrifuger* generally used the least memory, and *Metabuli* always needed substantially more memory than other tools (Fig. 4B).

**Figure 4.**
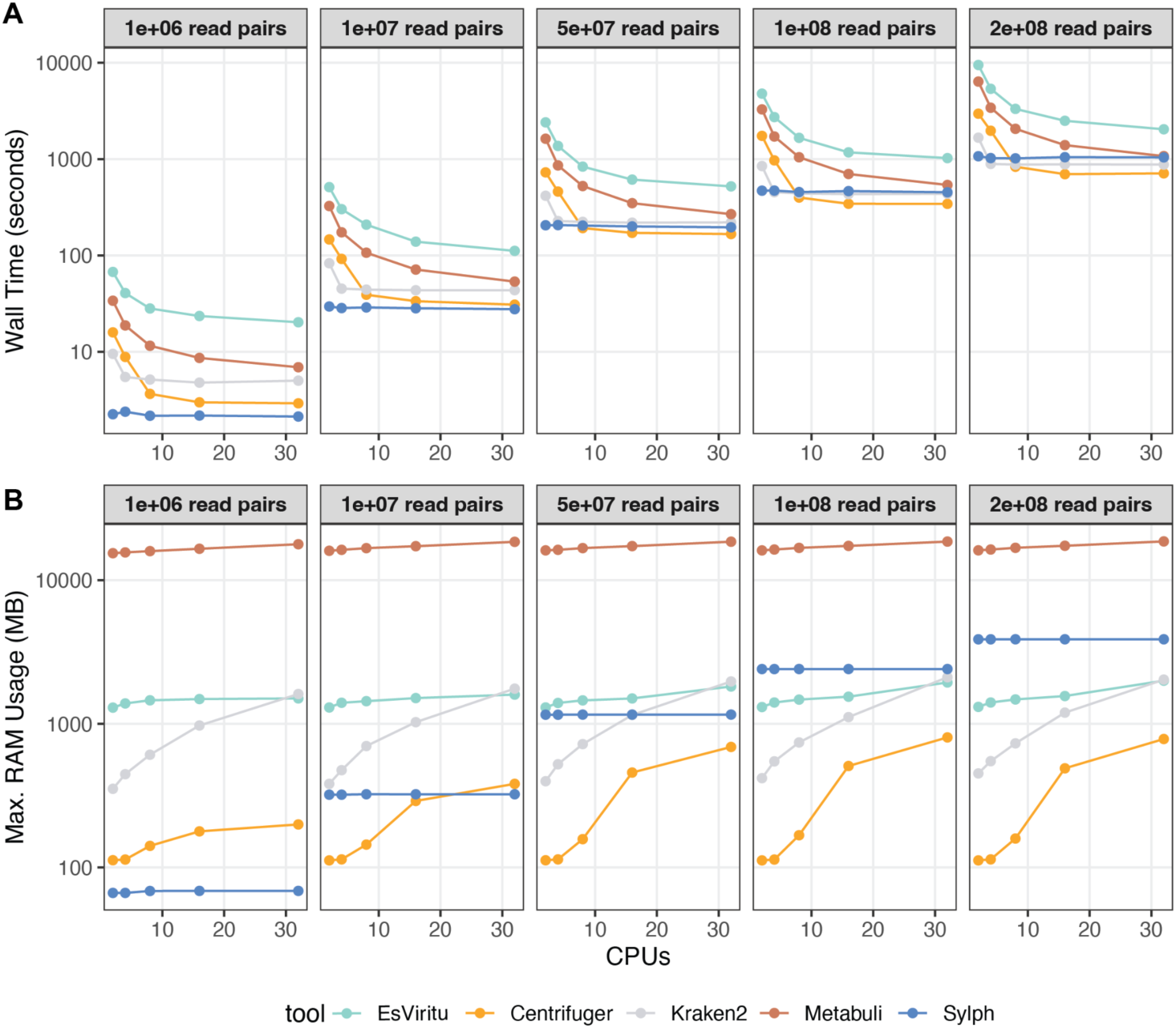
Computational Resource Scaling and Performance. (A) Wall time (y-axis) for each tool with increasing CPUs (x-axis) across sequencing read count. (B) Maximum RAM usage (y-axis) for each tool with increasing CPUs (x-axis) across sequencing read count.

## Discussion

At time of writing, *EsViritu* is already in use by many scientific groups around the world, especially for wastewater-based pathogen monitoring^34–37^, and the software has over 6,600 downloads on the Bioconda package manager. As unbiased or agnostic virus metagenomic sequencing continues to grow, we anticipate that this software will be key infrastructure in scientific and public-health workflows. The benchmarks reported here validate this potential.

*EsViritu* outperforms the leading metagenomic profilers for virus pathogen detection and quantification. Across many tools and parameter configurations while using the same Virus Pathogen Database, *EsViritu* is the most accurate at reconstructing community composition, either from a detection perspective (F1 score) or an abundance accuracy (Bray-Curtis dissimilarity, Pearson, and Spearman correlations to ground truth) (Fig. 3B, Fig. S2). This includes effective classification for simulated genomes at only 80%-85% similarity to nearest known reference. These mock-community results demonstrate that *EsViritu* is a superior choice for samples likely containing multiple viruses, where the goal is to reconstruct the composition and relative abundance of a whole viral community.

The positive and negative control results suggest that *EsViritu* is both sensitive and specific (Fig. 2). Therefore, this tool is equally suited to bulk sequencing (e.g. identifying a handful of rare virus reads in a clinical specimen without raising false alarms) or virus-enriched environmental samples, where virus-derived reads need to be properly classified even if they diverge from known genomes. A single tool that serves both regimes improves virome surveillance at a time when the importance of the virome is being recognized.

*EsViritu’s* advantage in virome profiling can be understood at the algorithm level. Other profilers tend to over-classify reads, i.e. assigning reads with confidence to species they cannot truly resolve and often assigning reads from one singular genome to various related taxa. This behavior explains the much of the pattern we observed regarding F1 scores and Bray-Curtis dissimilarity in mock communities (Fig. 3). By resolving read sharing among related references before committing to a call, and by refining taxonomy to the rank the data support, *EsViritu*’s algorithm avoids these common missteps (Fig. 3C).

We distribute *EsViritu* with the Virus Pathogen Database, which combines the comprehensiveness of nearly 20,000 publicly-available virus genomes across 63 families, with the added value of database-side host and vector contamination, genome clustering, and low-complexity masking (Methods). Despite the comprehensive diversity, *EsViritu* with the Virus Pathogen Database can be readily run on a couple CPUs and GB of RAM (Fig. 4). However, users can use the a few simple commands to extend or completely replace this database with other sequences, allowing for the algorithm and software to extend to custom use cases.

*EsViritu* also returns a consensus genome for each detection and reports its ANI to the nearest reference, and the reconstruction closely tracks ground truth (Fig. S4). This feature helps researchers verify results and conduct downstream genomic and phylogenetic analyses. The genome reconstructions also let analysts distinguish between similar strains and provide the sequence-level evidence that public-health and biosecurity decisions require. *EsViritu* additionally collates run- and project-level information in an interactive HTML report. Ultimately, a tool that is both highly sensitive and highly specific while returning multiple useful outputs is well-positioned to serve the pathogen monitoring community.

We also wish to flag several caveats regarding these conclusions. The *divergence-capping* metric reflects family-dependent viral species cutoffs and is a documented heuristic, so we reported strict and capped results. Further, novel genomic recombination and reassortment regularly occur in the virus world, but our benchmarks do not test the tools’ robustness to this phenomenon. Finally, our group’s extensive experience with *EsViritu* suggests that it can make mistakes, including classification of virus-derived vector sequences and over-confident calls resulting from using an incomplete genome database. Therefore, occasional manual validation may be necessary. None of these caveats alters the central finding that *EsViritu* is a valuable tool for analysis of virome data, which we show across positive controls, negative controls, and mock communities.

## Methods

### *EsViritu* algorithms and code

First, reads are aligned to the Virus Pathogen Database using minimap2^38^ with the following per-read filters: alignment fraction >= 90%, average nucleotide identity >= 80%, alignment length >= 100 nt. Secondary alignments are kept for alignment scores >= 0.9 of primary. Then, the most appropriate reference genomes for the sample’s data are determined through a greedy clustering algorithm based on read-sharing. This algorithm is “assembly-aware” in that multi-segmented viruses are quantified as a single assembly during clustering. The algorithm follows:

### Edge definition (graph *G*)

An undirected edge exists between assemblies *a* and *b* iff:

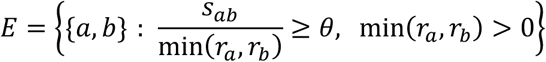

Neighborhood: *N*(*a*) = {*b*: {*a*, *b*} ∈ *E*}

### Ranking key

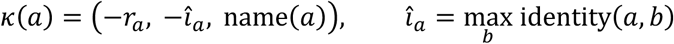

### Greedy assignment (repeat until *U* = ⌀)

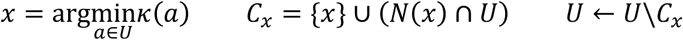

where *r_a_* = total reads, *s_ab_* = shared reads, *θ* = threshold (0.33), *U* = unassigned assemblies.

The read alignment-clustering steps are repeated an additional time. Then, a final minimap2 alignment is conducted, and genomes are reconstructed based on read sequences (i.e. reference-guided assembly). Coverage metrics and Pi (average nucleotide diversity per site) are calculated. Reconstructed genome records are aligned back to the Virus Pathogen Database, and the lowest common ancestor (i.e. deepest taxon with the unambiguously best alignment score) is used to define the taxonomy of the record. Identity thresholds are then used to roll back taxonomy by collapsing subspecies to the species (<= 95% ANI) or genus (<= 90% ANI) level if appropriate.

Next, RPKMF (Reads per Kilobase per Million Filtered reads; “filtered reads” meaning total reads passing quality filtering, whether assigned to a taxon or not) is calculated at the assembly level. Then, a taxonomic profile is made based on the taxonomic assignments and abundances of all assemblies.

Finally, a custom HTML report is prepared for interactive visual inspection of all records classified by EsViritu.

Additional utility scripts are available for custom database creation or supplementation and multi-sample output collation.

The main functions of *EsViritu* are written in Python (v3.10 or greater) and depend on the following software libraries: pysam, polars, pyyaml, numpy. The HTML report generation scripts are written in R and depend on htmltools, reactable, reactablefmtr. The Sparklines function is vendored from dataui to avoid installation complexity. Minimap2 seeding/extension parameters are available for short-read “sr”, ONT “lr:hq”, and PacBio “map-hifi”, and sensitive “sense” (sr modified for k=11 and w=10 for more aggressive seeding).

*EsViritu* is available on Bioconda and the source code is maintained on GitHub at <u>cmmr/EsViritu</u>. Full documentation is available at https://esviritu.readthedocs.io.

### Virus Pathogen Database

Virus Pathogen Database is a manually-curated and opinionated database of virus genome sequences and sequence-level metadata for taxonomy and sequence description. In brief, sequences and metadata were downloaded from NCBI Virus for the 63 virus families in which there is plausible evidence of human, animal, or plant infection. These were filtered for 1) “complete” labels or near-complete length and appropriate number of segments, 2) comprehensive taxonomy (e.g. sequences defined to the species level).

Then, all sequences per-family were compared all-vs-all using BLASTN^39^ and clusters were generated as defined by 95% ANI and 85% AF. Exemplar genomes were chosen by 1) RefSeq record then 2) longest sequence. There are several exceptions to the clustering for important pathogens which were clustered within the genotype (subspecies) level to preserve important sequence diversity: SARS-CoV-2, norovirus, monkeypox virus, rotavirus A, measles virus, dengue virus, influenza A virus. Norovirus genotypes were defined with Norovirus Genotyping Tool (v2.0)^40^, and rotavirus A genotypes were defined with <u>Rotavirus A Genotyping Tool</u> (v0.1).

Next, low complexity repeats were N-masked with bbtools^41^ bbmask on default settings. Genomes were aligned with minimap2 to human (GCF_000001405.40), dog (GCF_011100685.1), mouse (GCF_000001635.27), cow (GCF_002263795.3), macaque (GCF_003339765.1), chicken (GCF_016699485.2), cat (GCF_018350175.1), and rat (GCF_036323735.1) genomes. We also aligned the genomes to the UniVec vector database (NCBI) excluding SV40 virus derived sequences. Alignments of 80 nt or more to these genomes/vectors were N-masked. All exemplar genomes were then combined to generate the FASTA format sequence records. Taxonomic hierarchy labels were determined using NCBI taxids and pytaxonkit^42^ and added to other metadata distributed as a TSV file. At time of writing, the most recent version is v3.2.4^23^, published November 15, 2025 on Zenodo.

### Tools and parameters

We compared *EsViritu* v1.3.1 against *Kraken2* v2.17.1^19^, *Centrifuger* v1.1.1-r309^26^, *Metabuli* v1.2.0^27^, and *Sylph* v0.9.0^28^. All tools were able to use identical genome sequences from the Virus Pathogen Database and the same NCBI TaxDump file to create the tool specific database file. Each tool was run across a grid of its sensitivity- and specificity-governing parameters. All tools queried the same *EsViritu* Virus Pathogen Database, so reference content was held constant across the comparison. *EsViritu* was run in both its sensitive “*sense*” and short-read “*sr*” configurations. The Virus Pathogen Database v3.2.4 was used with one important modification: To allow cross-tool compatibility, the taxonomic hierarchy labels were pulled exclusively from the NCBI TaxDump from 2026-06-01, erasing any custom subspecies labels (e.g. for norovirus) that the standard v3.2.4 uses. This modified version “v3.2.4b” is available on Zenodo (10.5281/zenodo.23043160).

### Positive controls

Synthetic read sets were generated from seven reference genomes spanning 75.9% to 95.6% ANI to their closest database reference (Table S25), at depths of 1, 10, 100, and 1,000 reads with ten independent permutations per depth. A detection was scored when a tool reported the corresponding family. Detection was modeled with Firth-penalized logistic regression^43^ (detected ∼ tool x depth + genome, *EsViritu* as reference, genome as a fixed block) to handle complete separation for the most and least sensitive tools. Tool-versus-*EsViritu* contrasts used Benjamini-Hochberg correction^44^. LD50, the depth giving 50% detection, was estimated per genome and plotted against ANI; because ANI is confounded with genome (one genome per ANI level), divergence is treated descriptively rather than as a fitted coefficient. For this experiment, reads were synthesized with bbtools randomreads.sh.

### Negative controls

Pathogen-free samples were drawn from seven sources spanning host and synthetic backgrounds (human genome mock reads, mouse RNA) and environmental backgrounds (anaerobic digester, freshwater, marine water, hot springs, soil) (Table S16). False-positive read percentage and false-species counts were compared across tools using Friedman tests^45^ with post-hoc Wilcoxon tests (Benjamini-Hochberg corrected) and Kendall’s W effect sizes.

### Mock communities

Six mock communities were built in two designs: three communities of 90 viral genomes drawn randomly from the Virus Pathogen Database, and three communities of 60 genomes drawn with no more than one assembly per family. In all communities the genomes were mutated with a sequence simulator to a ladder of 80%, 85%, 90%, 95%, 98%, and 100% ANI to their references. A reproducible pipeline for this is available on GitHub at <u>mtisza1/mut_virome</u>. Reads were synthesized with InSilicoSeq, using log-normal distributions and Novaseq error model. 10,000 read pairs were synthesized for viruses, then a background of 100 random bacterial genomes plus the human genome was added at 50,000,000 reads (50,010,000 total reads/mock community). Mock Community read tables can be found in the Supplemental Data Zenodo repository (10.5281/zenodo.23043160).

Profiles were scored against ground truth at every rank for detection (F1) and abundance (Bray-Curtis dissimilarity^46^, Pearson correlation), with profiles normalized to relative abundance per tool. Abundance calibration used Lin’s concordance correlation coefficient^47^, computed in a pooled stratum (all detected genomes) and a resolvable stratum (ANI at least 95%, single-clade). Profiles were additionally compared to ground truth under strict (provenance-lineage) and divergence-capped truth, the latter using an ANI-to-rank cap (98%-100%: subspecies, 90%-95% species, 80%-85% genus); both modes are reported.

*Sylph* reports a k-mer-containment sequence-abundance value and was not readily comparable to other abundance metrics. All six communities were analyzed together. All analyses were performed in R.

### Genome Reconstructions

Genome-reconstruction accuracy was assessed in two ways. First, by comparing the *EsViritu*-estimated ANI of each consensus reconstruction to the ground-truth ANI of the source genome. Second, *EsViritu*-generated reconstructions were compared to the mutated genomes used as templates for the synthetic reads in the mock communities with BLASTN with default parameters, and non-N positions (i.e. canonical bases) were compared to determine total mismatches. We note that these mock communities often had viruses represented by only a handful of reads, and genome reconstructions are more accurate at higher coverage with lots of read support data.

### Code and data availability

EsViritu is available on <u>Bioconda</u> and the source code is on GitHub at <u>cmmr/EsViritu</u>, with full documentation on a <u>readthedocs</u> website. The Virus Pathogen Database v3.2.4 (doi 10.5281/zenodo.17716199), was published November 15, 2025 on <u>Zenodo</u>.

The processing and analysis repo for the benchmarks are maintained on <u>GitHub</u> with stable release <u>2026-09-29a</u>. The repo for generating synthetic mutations with random Virus Genome Database, *mut_virome* is available <u>on GitHub</u> with stable <u>release v0.1.0</u>.

All data to reproduce the analyses have been uploaded to Zenodo (10.5281/zenodo.23043160).

## Acknowledgements

We thank Adrien Assie for illustrating the logo for *EsViritu*. We acknowledge Anil Surathu, Lander De Coninck, and Lisa Faye for bug fixes to the codebase. We also thank Alex Jaffe, Rose Kantor, Alessandro Zulli, Nitin Shukla, and Vikram Chan-Herur for feedback on *EsViritu*.

## References

1. Chiu, C. Y. & Miller, S. A. Clinical metagenomics. Nat. Rev. Genet. 20, 341–355 (2019).

2. Gu, W., Miller, S. & Chiu, C. Y. Clinical Metagenomic Next-Generation Sequencing for Pathogen Detection. Annu. Rev. Pathol. 14, 319–338 (2019).

3. Justen, L. J. et al. Deep untargeted wastewater metagenomic sequencing from sewersheds across the United States. Preprint at 10.64898/2026.03.05.26345726 (2026).

4. Kantor, R. S. & Jiang, M. Considerations and Opportunities for Probe Capture Enrichment Sequencing of Emerging Viruses from Wastewater. Environ. Sci. Technol. 58, 8161–8168 (2024).

5. Nabel, C. S. et al. Virome capture sequencing does not identify active viral infection in unicentric and idiopathic multicentric Castleman disease. PloS One 14, e0218660 (2019).

6. Mascarenhas, A. C. et al. Metagenomic sequencing identifies potential respiratory pathogens in PCR-negative subset of surveillance samples. Sci. Rep. 16, 9308 (2026).

7. Clark, J. R. et al. Wastewater pandemic preparedness: Toward an end-to-end pathogen monitoring program. Front. Public Health 11, 1137881 (2023).

8. Tisza, M. et al. Wastewater sequencing reveals community and variant dynamics of the collective human virome. Nat. Commun. 14, 6878 (2023).

9. Javornik Cregeen, S., et al. Sequencing-Based Detection of Measles in Wastewater: Texas, January 2025. Am. J. Public Health 115, 1115–1119 (2025).

10. Kelly, M. S. et al. Nasal microbiome and phageome profiles are associated with prospective respiratory viral infection risk in school-age children. J. Allergy Clin. Immunol. 158, 408–420 (2026).

11. Justen, L. J., Grimm, S. L., Esvelt, K. M. & Bradshaw, W. J. Indoor air sampling for detection of viral nucleic acids. J. Aerosol Sci. 187, 106549 (2025).

12. Briese, T. et al. Virome Capture Sequencing Enables Sensitive Viral Diagnosis and Comprehensive Virome Analysis. mBio 6, e01491–01415 (2015).

13. Rushford, C. et al. Untargeted longitudinal ultra deep metagenomic sequencing of wastewater provides a comprehensive readout of expected and unexpected viral pathogens. Preprint at 10.1101/2025.10.27.25338874 (2025).

14. Miller, S. et al. Laboratory validation of a clinical metagenomic sequencing assay for pathogen detection in cerebrospinal fluid. Genome Res. 29, 831–842 (2019).

15. de Vries, J. J. C. et al. Benchmark of thirteen bioinformatic pipelines for metagenomic virus diagnostics using datasets from clinical samples. J. Clin. Virol. Off. Publ. Pan Am. Soc. Clin. Virol. 141, 104908 (2021).

16. Wylie, T. N. & Wylie, K. M. ViroMatch: A Computational Pipeline for the Detection of Viral Sequences from Complex Metagenomic Data. Microbiol. Resour. Announc. 10, e01468–20 (2021).

17. Lu, D. et al. Simultaneous detection of pathogens and antimicrobial resistance genes with the open source, cloud-based, CZ ID platform. Genome Med. 17, 46 (2025).

18. Kalantar, K. L. et al. IDseq-An open source cloud-based pipeline and analysis service for metagenomic pathogen detection and monitoring. GigaScience 9, giaa111 (2020).

19. Wood, D. E., Lu, J. & Langmead, B. Improved metagenomic analysis with Kraken 2. Genome Biol. 20, 257 (2019).

20. Ye, S. H., Siddle, K. J., Park, D. J. & Sabeti, P. C. Benchmarking Metagenomics Tools for Taxonomic Classification. Cell 178, 779–794 (2019).

21. Gregori, J. et al. Viral quasispecies complexity measures. Virology 493, 227–237 (2016).

22. Huson, D. H., Auch, A. F., Qi, J. & Schuster, S. C. MEGAN analysis of metagenomic data. Genome Res. 17, 377–386 (2007).

23. Tisza, M. Virus Pathogen Database for EsViritu. Zenodo 10.5281/ZENODO.17716199 (2025).

24. Nasko, D. J., Koren, S., Phillippy, A. M. & Treangen, T. J. RefSeq database growth influences the accuracy of k-mer-based lowest common ancestor species identification. Genome Biol. 19, 165 (2018).

25. R Marcelino, V., Holmes, E. C. & Sorrell, T. C. The use of taxon-specific reference databases compromises metagenomic classification. BMC Genomics 21, 184 (2020).

26. Song, L. & Langmead, B. Centrifuger: lossless compression of microbial genomes for efficient and accurate metagenomic sequence classification. Genome Biol. 25, 106 (2024).

27. Kim, J. & Steinegger, M. Metabuli: sensitive and specific metagenomic classification via joint analysis of amino acid and DNA. Nat. Methods 21, 971–973 (2024).

28. Shaw, J. & Yu, Y. W. Rapid species-level metagenome profiling and containment estimation with sylph. Nat. Biotechnol. 43, 1348–1359 (2025).

29. Govender, K. N. & Eyre, D. W. Benchmarking taxonomic classifiers with Illumina and Nanopore sequence data for clinical metagenomic diagnostic applications. Microb. Genomics 8, mgen000886 (2022).

30. Yue, J.-X. & Liti, G. simuG: a general-purpose genome simulator. Bioinformatics 35, 4442–4444 (2019).

31. Gourlé, H., Karlsson-Lindsjö, O., Hayer, J. & Bongcam-Rudloff, E. Simulating Illumina metagenomic data with InSilicoSeq. Bioinformatics 35, 521–522 (2019).

32. Simmonds, P. et al. Virus taxonomy in the age of metagenomics. Nat. Rev. Microbiol. 15, 161–168 (2017).

33. Meyer, F. et al. Critical Assessment of Metagenome Interpretation: the second round of challenges. Nat. Methods 19, 429–440 (2022).

34. Wolfe, M. K. et al. Prospective metagenomic sequencing of wastewater across the United States yields robust viral enrichment and concordance with digital PCR measurements. Preprint at 10.64898/2026.05.07.26352651 (2026).

35. Shukla, N. et al. Longitudinal wastewater virome surveillance unveils untapped circulating viruses in the community. Npj Emerg. Contam. 2, 15 (2026).

36. Li, X. et al. Capture sequencing demonstrates promising public health potential for post-COVID wastewater surveillance: a comparative multi-technique and spatiotemporal analysis. Environ. Int. 209, 110146 (2026).

37. Karatas, M. et al. Untapped potential of wastewater for animal and potentially zoonotic virus surveillance: Pilot study to detect non-human animal viruses in urban settings. Environ. Int. 199, 109500 (2025).

38. Li, H. Minimap2: pairwise alignment for nucleotide sequences. Bioinformatics 34, 3094–3100 (2018).

39. Camacho, C. et al. BLAST+: architecture and applications. BMC Bioinformatics 10, 421 (2009).

40. Kroneman, A. et al. An automated genotyping tool for enteroviruses and noroviruses. J. Clin. Virol. Off. Publ. Pan Am. Soc. Clin. Virol. 51, 121–125 (2011).

41. Bushnell, B. & Villada, J. C. Deployable high-fidelity metagenome binning at scale with QuickBin. Commun. Biol. 9, 1135 (2026).

42. Shen, W. & Ren, H. TaxonKit: A practical and efficient NCBI taxonomy toolkit. J. Genet. Genomics Yi Chuan Xue Bao 48, 844–850 (2021).

43. Firth, D. Bias reduction of maximum likelihood estimates. Biometrika 80, 27–38 (1993).

44. Benjamini, Y. & Hochberg, Y. Controlling the False Discovery Rate: A Practical and Powerful Approach to Multiple Testing. J. R. Stat. Soc. Ser. B Stat. Methodol. 57, 289–300 (1995).

45. Friedman, M. The Use of Ranks to Avoid the Assumption of Normality Implicit in the Analysis of Variance. J. Am. Stat. Assoc. 32, 675–701 (1937).

46. Bray, J. R. & Curtis, J. T. An Ordination of the Upland Forest Communities of Southern Wisconsin. Ecol. Monogr. 27, 325–349 (1957).

47. Lin, L. I. A concordance correlation coefficient to evaluate reproducibility. Biometrics 45, 255–268 (1989).

